# Toxicity of microplastics and plastic additive co-exposure in liver Disse organoids from healthy donors and patient-derived induced pluripotent stem cells

**DOI:** 10.1101/2022.09.12.506301

**Authors:** Shaojun Liang, Yixue Luo, Jun Yi, Lu Feng, Mingen Xu, Rui Yao

**Author notes:** Corresponding author (Rui Yao).

## Abstract

The ubiquitous microplastics (MPs) and plastic additives in the environment usually form complexes, enter human blood circulation, and increase the risk of steatohepatitis. The liver Disse space plays a vital role in corresponding hepatic pathological processes. However, due to the limited understanding of the regulatory cues in multilineage maturation, the generation of large-scale Disse-like organoids (DOs) mirroring the comprehensive toxicity responses of MPs is challenging. Here, using human-induced pluripotent stem cells (hiPSCs), we biofabricated healthy donors and patient-derived DOs containing hepatocytes, endothelial cells, and hepatic stellate cells, resembling the features of Disse space. These organoids revealed that polystyrene MPs preferentially entered endothelial cells and then dispersed throughout the organoids, similar to reported studies in zebrafish. Co-exposure to MPs and tetrabromobisphenol A (TBBPA), a common plastic additive, showed enhanced accumulation of contamination in the organoids. We also biofabricated alcoholic liver disease (ALD) patient-derived DOs representing the specific disease transcriptional profiles. We found that co-exposure to MPs and TBBPA at environmental-related dosages significantly elevated the pathological transcriptional expression and biochemical profiles in patient-derived DOs but not in healthy organoids, suggesting that both hereditary factors and pollutants contribute to susceptibility to environmental toxicants. This study exemplified the value of biofabricated hiPSC-derived organoids in environmental toxicology and offered a powerful strategy for personalized toxicology evaluation.

## 1. Introduction

Daily environmental exposure to MPs and their potential effects on human health have raised public concerns in recent years. MPs (<10 µm) were detected in the blood of healthy individuals^1^ and the placenta^2^ and accumulated in human cirrhotic liver tissues^3^. Besides, MPs as vectors transfer plastic additives and toxicants in the environment into the body, thus causing combined toxicity^4^. Among these toxicants, TBBPA is a commonly used brominated flame retardant and potential endocrine disruptor ^5^. Co-exposure to MPs and TBBPA disturbed hepatic redox status more severely than single exposure in zebrafish^6^. Altered redox balance leads to liver cirrhosis, including activation of hepatic stellate cells (HSCs) that reside in the liver Disse space, where hepatocytes and hepatic sinusoidal endothelial cells regulate the quiescent and activated states of HSCs^7^. Compared to monocultures, cocultures of liver stem cells (HepaRG), endothelial cells (HUVECs), and HSCs (LX-2) dispersed in bioprinting structures better recapitulated the critical steps of fibrogenesis^8^. However, due to the lack of *in vitro* models mimicking close cell-cell interactions in the heterogeneous human Disse space, the hepatotoxicity mechanisms of MPs and plastic additives are open questions, particularly in environmental-related concentrations.

Multicellular organoids developed from human pluripotent stem cells can recapitulate tissue structure and multifaceted functions by re-establishing heterogeneous cell-cell interactions and are regarded as advanced alternatives in disease modeling, drug screening and autologous therapy^9^. Recently, a milestone report on human embryonic stem cell-derived hepatocyte organoids revealed the toxicology of 1 µm polystyrene MPs by disrupting liver lipid metabolism^10^. Human induced pluripotent stem cells (hiPSCs) carry genetic inheritance of individuals^9^; thus, hiPSC-derived organoids are potent in personalized environmental toxicology. However, due to the limited understanding of the regulatory cues in the multilineage maturation of hepatic/endothelial/hepatic stellate cells and suitable mass-production methods, hiPSC-derived organoids have not yet been applied in environmental toxicity studies.

Here, we mass-produced multicellular DOs based on healthy donors, patient-derived hiPSCs, and electro-assisted inkjet bioprinting. The chemically defined alginate/laminin bioinks and 3D microenvironment of microspheres facilitated multicellular crosstalk and mutually enhanced the maturation of hiPSC-derived progenitor cells, thus generating DOs mimicking Disse space features. MPs (1 µm) crossed the barrier of the hydrogel matrix and were preferentially taken up by endothelial cells and then dispersed throughout the whole organoid. Co-exposure to MPs and TBBPA enhanced pollution accumulation at the tested time points. ALD patient-derived hiPSC-formed DOs representing the specific disease transcriptional profiles efficiently, suggesting the robustness of the bioprinting organoid approach. Transcriptional and biochemical evaluations suggested that co-exposure to MPs and TBBPA severely impacted patient-derived DOs over healthy donor organoids **(Fig. 1A)**. This study takes us one step further into the generation of hiPSC-based bioprinting organoids and personalized environmental toxicology paradigms.

**Fig 1.**
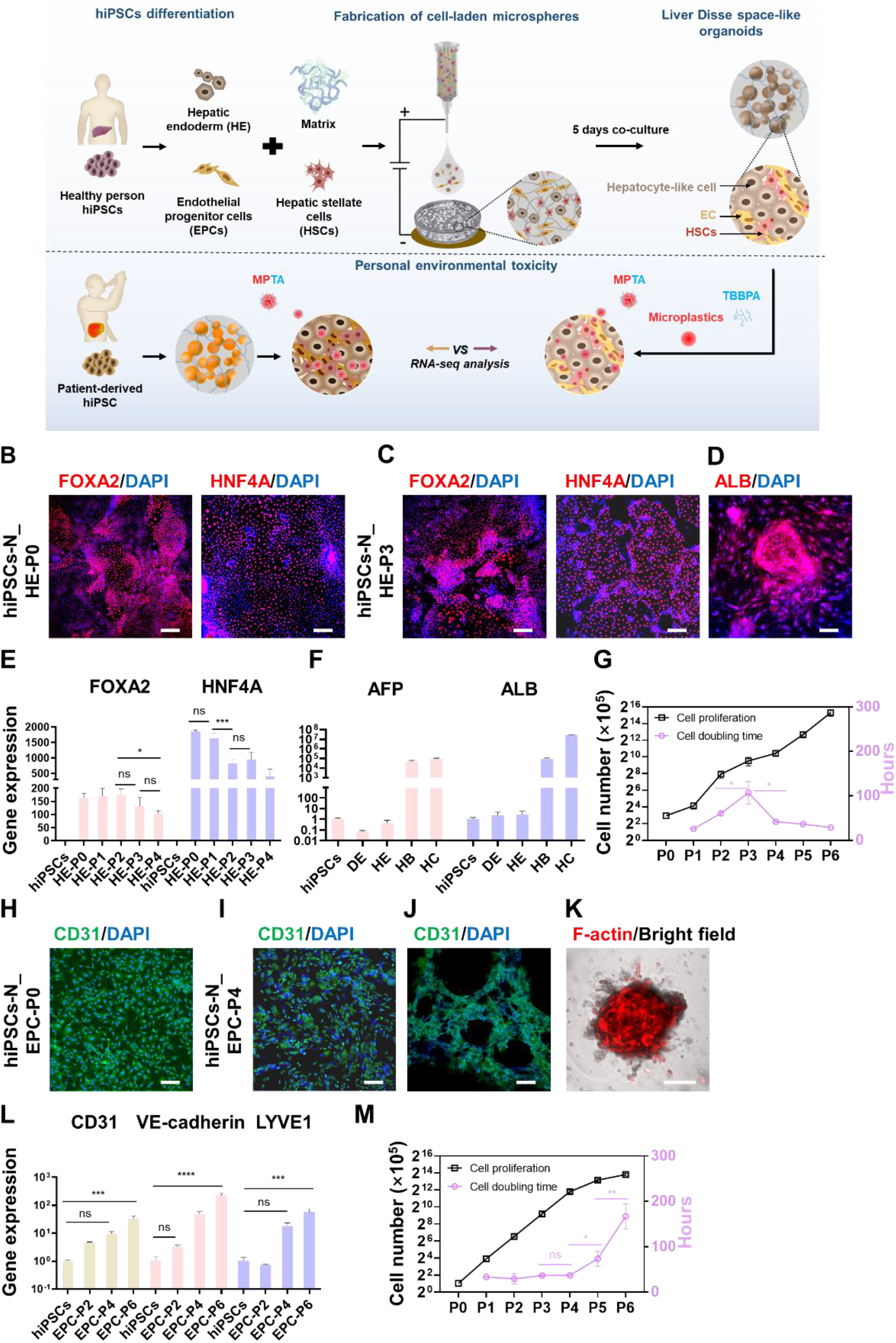
Generation of hepatic endoderm and endothelial-like cells from hiPSCs-N. (A) Schematic illustration of the fabrication process of hiPSC-derived DOs. Immunofluorescence staining (IF) of hepatic endoderm (HE) markers (FOXA2 and HNF4A) in hiPSCs-N_HE-P0 (B) and hiPSCs-N_HE-P3 (C). (D) IF of ALB in hiPSC-N_HE-P3-derived hepatocyte-like cells. Scale bars in B-D, 200 µm. qPCR of HE markers in HE_P0 to P4 (E), hepatocyte markers in DE, HE, HB, and HC (F), normalized to GAPDH and hiPSCs. G) The cell proliferation (black curve) and cell number doubling time (pink curve) of hiPSCs-N_HE. IF of the endothelial marker CD31 in hiPSCs-N_EPC-P0 (H) and hiPSCs-N_EPC-P4 (I). (J) Maximal projection of the vascularization of hiPSCs-N_EPC-P4 in the Matrigel. Scale bars in H-J, 200 µm. (K) Maximal projection of F-actin staining and brightfield image of hiPSC-N_EPC-P4-derived organoids. Scale bar, 100 µm. (L) qPCR of endothelial markers in hiPSCs-N_EPC-P0 to P4, normalized to GAPDH and hiPSCs. (M) The cell proliferation (black curve) and cell number doubling time (pink curve) of hiPSCs-N_EPCs. The data in (E), (F), (G), (L), and (M) are presented as the mean ± s.d. Statistical significance was analyzed using one-way ANOVA with Tukey’s post-hoc test, \**p*<0.05, \*\**p*<0.01, \*\*\**p*<0.001, \*\*\*\**p*<0.0001, ns indicates no significant difference, n=3.

## Results and discussion

### 2. Fabrication of hiPSC-derived multicellular DOs

The Disse space harboring hepatocytes, sinusoid endothelial cells, and hepatic stellate cells plays a crucial role in the liver response to toxic substances^11^. We sought to tackle the challenges of large-scale biofabrication of biomimetic DOs from two main aspects: (1) high-quality and sufficient amounts of hiPSC-derived progenitor cells as cell source and (2) biofabrication methods considering both the survival of hiPSC-derived cells and multilineage maturation.

#### 2.1. Specification of hiPSCs into progenitor cells

In this study, hepatic endoderm cells (HEs) and endothelial progenitor cells (EPCs) were derived from hiPSCs and planar differentiation and expansion. High-quality HEs and EPCs with desired cell identity and suitable proliferative capacity were selected for subsequent biofabrication and organoid formation.

Healthy donor-derived hiPSCs (hiPSCs-N) were directed into FOXA2+/HNF4A+ HE (counted as passage 0, HE-P0) (Fig. 1B, 1E). After planar expansion for 3 passages, HE-P3 maintained the positive cell identity markers (Fig. 1C, 1 E), while more than 3 passages showed declined marker expression and loss of cellular morphology (Fig. 1E, 1G). Given that HE-P3 showed the potential of hepatocyte specification with positive ALB/AFP expression (Fig. 1D, 1F), hiPSC-N-derived HE-P3 was used for DO biofabrication.

hiPSCs-N were also directed into CD31+ EPCs (Fig. 1H, 1 L). Propagation in planner conditions enhanced EPC maturation with upregulation of transcriptional endothelial markers (CD31, VE-cadherin, and LYVE1) (Fig. 1I, 1L), and thus EPC-P4 cells were capable of vascularization (Fig. 1J, 1K, Fig. S1, S video 1). Given that EPCs from P5 showed a decreased cell doubling time (Fig. 1M), hiPSC-N-derived EPC-P4 cells were used for DO biofabrication.

#### 2.2 Biofabrication of Disse space-like organoids

Given that hiPSC-derived cells are sensitive to mechanical and chemical stimulation, the mechanical force of 3D printing and the chemical properties of bioinks significantly constrict the generation of liver organoids from hiPSC-derived monodisperse cells^12^. Our previous study found that electroassisted inkjet printing of alginate/gelatin microspheres offered a physical and biochemical environment that facilitated uniform aggregation of stem cells and stemness maintenance^13^. Because laminin 511 promotes organoid survival and liver development^14,15^, we optimized the chemically defined bioink of alginate and laminin 511, replacing gelatin. With printing technology and bioinks, we massively produced hydrogel microspheres (1×10^7^ cell-laden microspheres per milliliter) with diameters of 498±57 µm (Fig. S2A). Within the microspheres, hiPSC-N-derived HE and EPC cells, together with the human HSC cell line LX-2, were assembled into organoids (N_DOs) with diameters of 32-44 µm (Fig. 2A, S2B). The viability of the postprinted cells was higher than 90% and almost recovered after five days of culture (Fig. 2A). By extrusion bioprinting, the viability of the hiPSC-derived single-cell dispersion was only 60%^12^. Taken together, we established an engineering platform suitable for printing monodisperse hiPSC-derived cells with improved cell viability.

**Fig 2.**
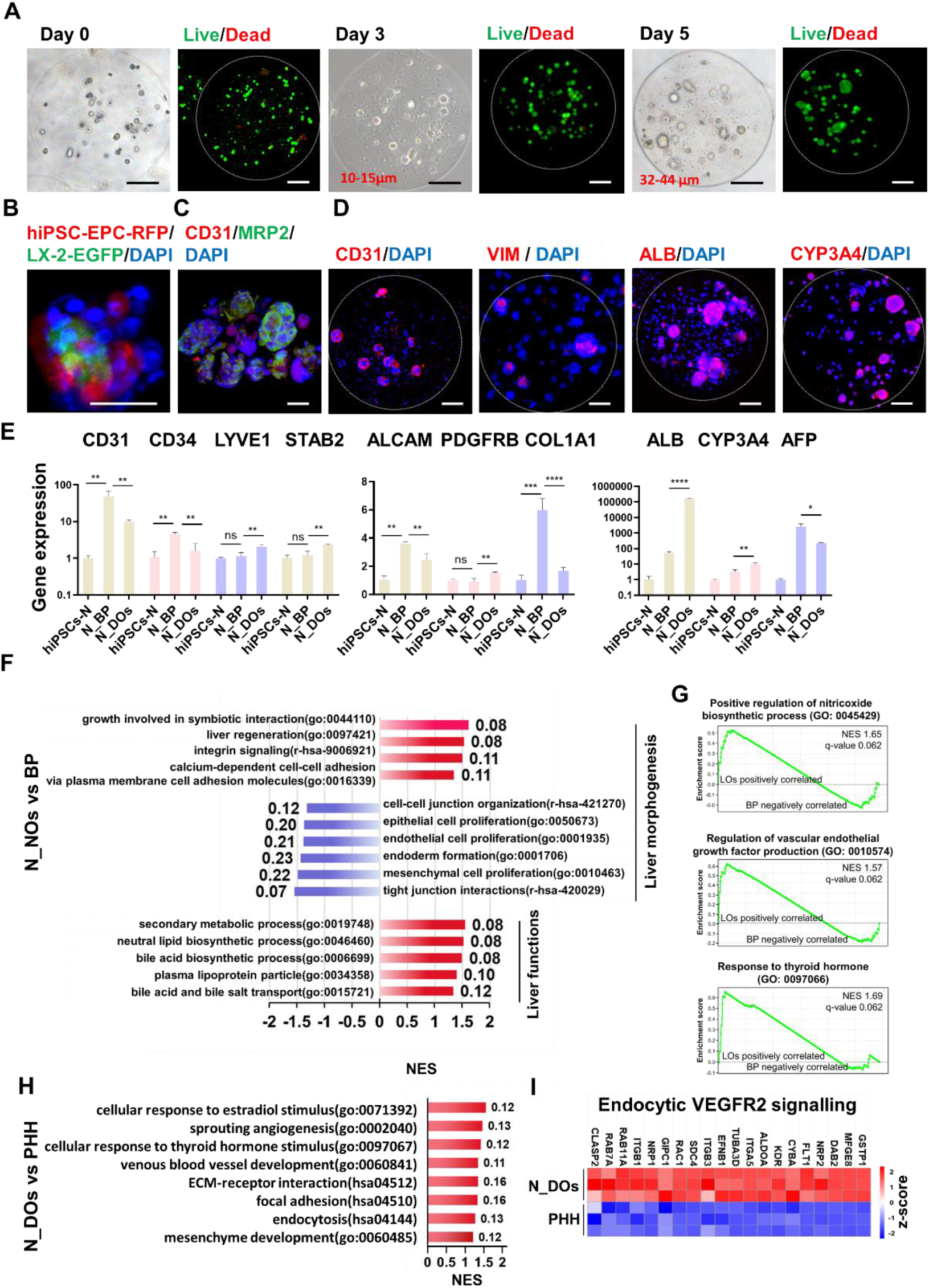
Fabrication of hiPSC-N-derived DOs. (A) Optical images and maximal projection of live/dead images of postprinted cell-laden microspheres on days 0, 3, and 5. Scale bars, 100 µm. (B) Maximal projection of the immunostaining image of N_DOs composed of hiPSCs-EPC-RFP, LX2-EGFP, and DAPI. Scale bar, 25 µm. (C) The maximal projection of in situ immunostaining images of hepatocyte markers (ALB and CYP3A4), endothelial marker (CD31), and HSC marker (VIM) in N_DOs within the microspheres. The dashed circles in (A) and (C) mark the outline of the microspheres. (D) The maximal projection of immunostaining images of CD31, MRP2, and DAPI in N_DOs harvested from microspheres. Scale bars in C-D, 50 µm. (E) qPCR of endothelial markers (CD31, CD34, LYVE1, STAB2), HSC markers (ALCAM, PDGFRB, COL1A1), hepatocytes (ALB, CYP3A4, AFP), normalized to GAPDH and hiPSCs. The data in (E) are three technical replicates of RNA pooled from three baths of organoids. Data are presented as the mean ± s.d. Statistical significance was analyzed using one-way ANOVA with Tukey’s post-hoc test, \**p*<0.05, \*\**p*<0.01, \*\*\**p*<0.001, \*\*\*\**p*<0.0001. (F) GSEA of GO terms based on the differential expression between N_DOs and BP. (H) GSEA of GO/Reactome terms based on the differential expression between N_DOs and PHHs. In F-H, the X axes represent the NES (normalized enrichment score); the numbers next to the columns represent the FDR q-value. (G) Representative enplots of GSEA for the comparison of N_DOs and BP. (I) Heatmaps of gene expression in N_DOs and PHHs; the bar stands for z score values, n=3.

Then, we used fluorescence imaging and qRT‒PCR to characterize the multicellular organoid assembly. hiPSC-EPC-RFP, LX-2-EGFP, and hiPSC-HE (stained with Hoechst 33342) were self-assembled into N_DOs in the hydrogel microspheres. CD31^+^ EPCs closely interacted with MRP2^+^ hepatocytes in N_DOs, incorporating CD31^+^ EPC, VIM+HSC, and ALB+/CYP3A4 + hepatocytes in the view of in situ microspheres, suggesting that N_DOs resembled the multicellular interaction of Disse space (Fig. 2B, 2C, 2D). Compared to the cell mixture before printing (BP), N_DOs exhibited the upregulation of LSEC marker genes (LYVE1 and STAB2) coupled with decreased negative genes (CD31 and CD34), upregulation of HSC marker genes (ALCAM, VIM, PDGFRB) associated with decreased HSC activation-related genes (COL1A1), and upregulation of hepatocyte mature genes (ALB, CYP3A4) connected with fetal gene (AFP) (Fig. 2E). Therefore, biomimetic cell composite and multicellular interactions within hydrogel microspheres contributed to Disse space-specific phenotypes.

To reveal the liver morphogenesis and functions of N_DOs, we conducted RNA-seq and GSEA by comparing N_DOs with BP and PHH, the standard for in vitro hepatotoxicology. Compared to BP from planner culture conditions, N_DOs showed upregulation of morphogenesis of multicellular organoids, probably by downregulating cell proliferation and tight junctions and by upregulating motility, integrin signaling, and calcium-dependent cell‒cell interactions (Fig. 2F). The three cells in the organoids were synergistically developed and mature (Fig. 2F), including phase Ⅰ/Ⅱ metabolism, hormone response, vascular endothelial growth factor production, and nitric oxide biosynthetic processes (Fig. 2G, Fig. S2B). Compared to PHH, N_DOs advanced in morphogenesis and function-related terms such as angiogenesis, vessel development, ECM-receptor interaction, cellular response to thyroid hormone, estradiol stimuli and endocytosis (Fig. 2H). The upregulated endocytic VEGFR2 signaling highlighted the liver endocytic capacity of N_DOs more than PHHs (Fig. 2I). These results indicate that N_DOs are more representative in the context of morphogenesis and functions in response to toxicants than PHHs.

Taken together, we developed a bioprinting workflow for the massive fabrication of hiPSC-derived multicellular liver organoids from monodisperse hiPSC-derived cells. The biomimetic cell composites and multicellular interaction enabled N_DOs to resemble 3D morphogenesis and diverse liver functions, thus laying the foundation to mirror the multifaceted hepatotoxicity of exogenous toxicants.

### 3. Diffusion of the MPs and TBBPA complex in DOs

Diverse types and shapes of MPs, including fragments and microbeads, have been detected in human blood of healthy donors^1^ and patients’ liver samples^3^. In human cirrhosis liver tissue, the microbead MPs were made from polystyrene^3^. However, in liver tissue, whether MPs remain in the extracellular matrix or enter liver cells and the effect of TBBPA on MP accumulation are poorly understood. To investigate these questions, we observed the diffusion of fluorescently labeled spherical polystyrene MPs in the DOs within hydrogel microspheres and the effect of TBBPA at the environmental concentration on the accumulation of MPs in the DOs.

After 1800 s of incubation with 1 µm red fluorescence-labeled MPs, the fluorescence intensity became apparent in the organoid, whereas no fluorescence signaling was detected in empty microspheres, demonstrating that PS-MPs could cross the barrier of the extracellular matrix (ECM) into N_NOs (Fig. 3A, 3B, 3G).

**Fig 3.**
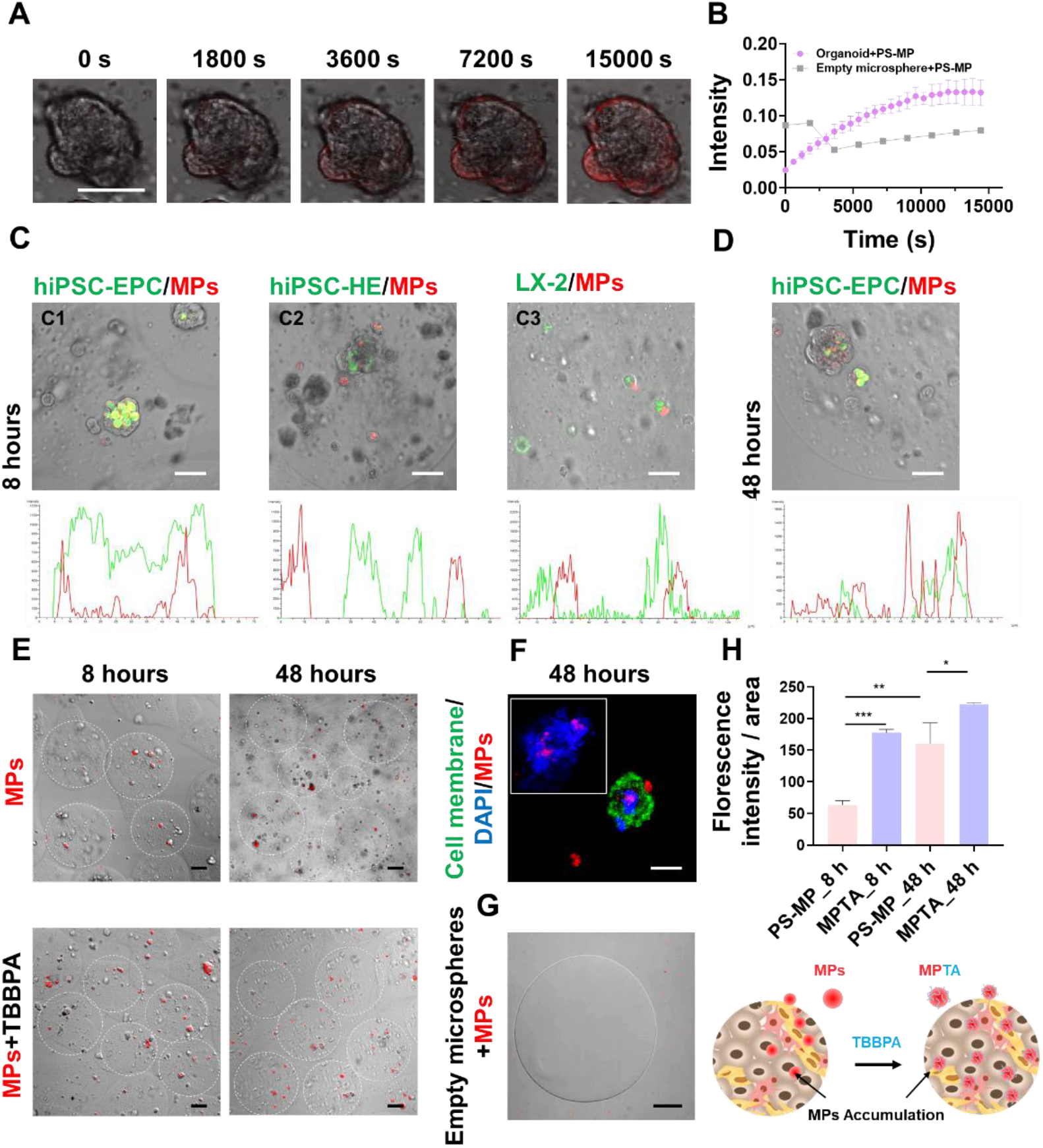
Diffusion of MPs in the DOs. (A) Time-lapse images of fluorescent images of MPs (red) merged with brightfield images of DOs. (B) The pink curve and blue curve depict the time-lapse fluorescence intensity of MPs in the organoid and an empty microsphere; the data are presented as the mean ± s.d. n=3. (C) and (D) The merged images of bright fields, red fluorescent MPs, and PKH67-labeled cell membranes of hiPSCs-N_EPC, HE, and LX-2, respectively. Scale bars in (A) and (C), 50 µm. The curves depict the localization of fluorescent signaling of MPs (red) and PKH67. (E) Merged images of red fluorescent MPs with brightfields of DO-laden microspheres. The dashed circles mark the outline of the microspheres. Scale bars, 100 µm. (H) Quantification of (E) was calculated as the fluorescence intensity of MPs divided by the microsphere area; over 20 microspheres were counted. The data in (H) are presented as the mean ± s.d. Statistical significance was analyzed using one-way ANOVA with Tukey’s post-hoc test, \**p*<0. 05, \*\**p*<0. 01, \*\*\**p*<0. 001, n=3. (F) MPs (red) penetrate the cell membrane labeled with PKH67 and nuclei labeled with DAPI. Scale bar, 10 µm. (G) The merged image of fluorescent MPs and the brightfield image of empty microspheres. Scale bar, 100 µm.

We labeled HE, EPC, and LX-2 cells with PKH67 membrane fluorescent dye to investigate whether MPs can be endocytosed into cells. After 8 hours of exposure, MPs entered the EPC more preferentially than hepatocyte-like cells (differentiated from HE) and LX-2 cells. MP signaling (red) entirely overlapped with EPC signaling (green) and dislocated from HE and LX-2 signaling (green) (Fig. 3C1-3). When extended to 48 hours, MP signaling appeared outside the region of hiPSC-N_EPC signaling, demonstrating the diffusion of MPs in N_DOs out of EPCs (Fig. 3D). Surprisingly, MPs were able to penetrate the cell membrane into the nucleus in the DOs, implying a disruption in genomic stability and gene transcriptional expression (Fig. 3E, 3F, 3H). Our finding was distinct from the study that 1 µm polystyrene MPs could enter planar cultured epithelial cells but not the nucleus region ^16^, probably due to the difference between 3D and 2D cultivation. With prolonged exposure, 1 µm polystyrene MPs accumulated in N_DOs, consistent with in vitro/vivo studies showing that polystyrene MPs emerged and accumulated in human hepatocytes and animal livers^4,17^.

Studies on toxicity interactions demonstrated that polystyrene MPs modified the biotransformation of TBBPA and aggravated its accumulation in vivo^18,19^. We found that TBBPA could aggravate the diffusion and accumulation of MPs in N_DOs (Fig. 3E, 3H), suggesting the mechanism of their synergistic toxicities^6^.

These results suggested that MPs could be preferentially endocytosed by endothelial cells and accumulated in the DOs. Due to the emergence of MPs in the nuclear region and aggravation of MP accumulation by TBBPA, the effects of the co-exposure of MPs and TBBPA on gene transcription need to be further evaluated.

### 4. The MP-TBBPA complex had little toxicity to healthy human-derived DOs

Based on human embryonic stem cell-derived hepatocyte organoids, polystyrene MPs caused cytotoxicity and disruption of gene expression and lipid metabolism, implicating the potential risks of liver steatosis, fibrosis and cancer^10^. However, since MPs are rarely detected in human liver tissue without disease, it is inconclusive whether MPs are safe for healthy individuals’ livers^3^. Based on healthy human hiPSC-derived Disse space-like N_DOs, from the perspective of multicellular interactions, we further investigated the cytotoxicity, transcriptomic and functional toxicities of spherical polystyrene MPs and TBBPA at environmentally relevant concentrations for 3 days. MPs were at 600 ng/mL, close to the concentration in bottled water of 656.8 ng/mL±632.9^20^; TBBPA was at 100 nM, slightly higher than 4.87 µg/L (equal to 9 nM) in lake water^21^; MPTA was at the combined dosages. (Fig. 4A).

**Fig 4.**
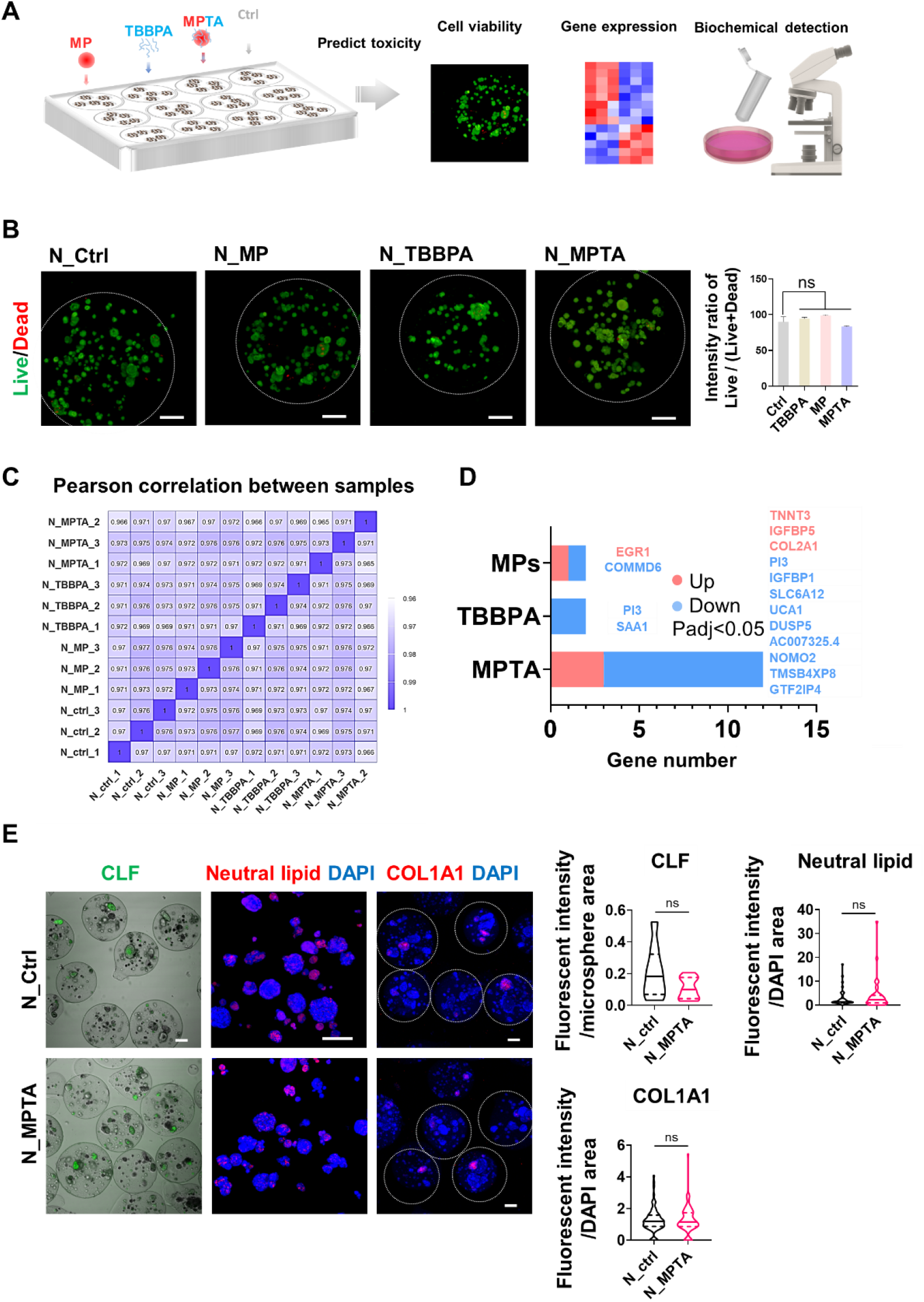
The effects of PS-MPs, TBBPA, and MPTA on hiPSC-N_DOs. (A) Schematic illustration of the toxicity assessment of PS-MPs, TBBPA, and MPTA with N_DOs. Maximal projection of the in situ staining of Live/Dead in N_DOs within microspheres treated with MPs, TBBPA, and MPTA for 3 days. Scale bars, 100 µm. The dashed circles mark the outline of the microspheres. The right panel is the ratio of Live/Dead calculated based on the three fields of Live/Dead staining. (C) The Pearson correlation heatmap of gene expression levels between all samples. (D) The parallel set depicts the relationships of the DEGs with GO terms based on differential expression between N_MPTA and N_ctrl. (E) Merged images of in situ staining of CLF merged with bright fields of N_DOs within microspheres; maximal projection of neutral lipid staining and in situ staining of COL1A1 and DAPI in N_DOs within the microspheres. Scale bars, 100 µm. The quantification of CLF, neutral lipids and COL1A1 based on the images is shown in the lower panels. The statistical significance was determined by Student’s T test; ns indicates no significant difference.

We used live/dead staining to detect cytotoxicity. The three groups of contaminants did not impact cell viability after 3 days of repeated exposure (Fig. 4B), consistent with most studies showing that MPs at concentrations below 1 µg/mL and TBBPA at nanomolar dosages rarely caused cytotoxicity^22,23^.

To reveal the adverse effects, we conducted RNA-seq analysis to compare N_MPTA, N_TBBPA, and N_MPTA to N_ctrl and found no differences in the Pearson correlation of transcriptomes within and between groups (values approximately 0.97) (Fig. 4C). MPs and TBBPA alone affected just 2 genes. MPTA disturbed 12 genes (Fig. 4D), among which the upregulated genes were IGF3P1, PI3, DUSP5, and SLC6A12, and the downregulated genes were TNNT3, IGF3P5, and COL2A1, which are related to insulin-like growth factor binding, endoplasmic reticulum, extractable matrix, myofilament, MAP kinase phosphatase activity, and transporter (Fig. 4D, S3). However, functional toxicity analysis by staining revealed that MPTA did not alter these gene-related liver functions, including bile acid transportation (CLF), neutral lipids, and COL1A1 accumulation in DOs (Fig. 4E).

Taken together, the hepatotoxicity of MPs, TBBPA, and MPTA at environmental concentrations was tolerable in N_DOs, suggesting that MPs and TBBPA are safe for healthy individuals. These results are distinct from the study mentioned above with human embryonic stem cell-derived hepatocyte organoids with dynamic exposure, which showed cytotoxicity, lipid metabolism, and related gene transcriptional toxicity at even lower concentrations of 250 ng/mL^10^. We reasoned that hydrogel encapsulation and multicellular constituents probably altered the toxic response, and second, static exposure in this study reduced the physical damage by particles.

### 5. DOs with ALD genetic backgrounds exhibited disease-related gene phenotypes

MPs were detected in the liver tissues of patients with chronic liver diseases, including ALD^3^. To investigate in depth the relationship between MPs/TBBPA and pathogenesis in patients, we adapted hiPSCs_AC19, derived from an ALD patient, to fabricate AC19_DOs and verify the representativeness of the genetic characteristics of ALD.

Briefly, hiPSC-AC19-derived AC19_HE-P3 and AC19_EPC-P4, together with LX-2, were printed and self-assembled into highly viable AC19_DOs (Fig. 5A, 5B, 5C). After coculture, multicellular AC19_DOs maturely expressed endothelial cell and hepatocyte markers (CD31/MRP2) and relevant genes (Fig. 5D, 5E).

**Fig 5.**
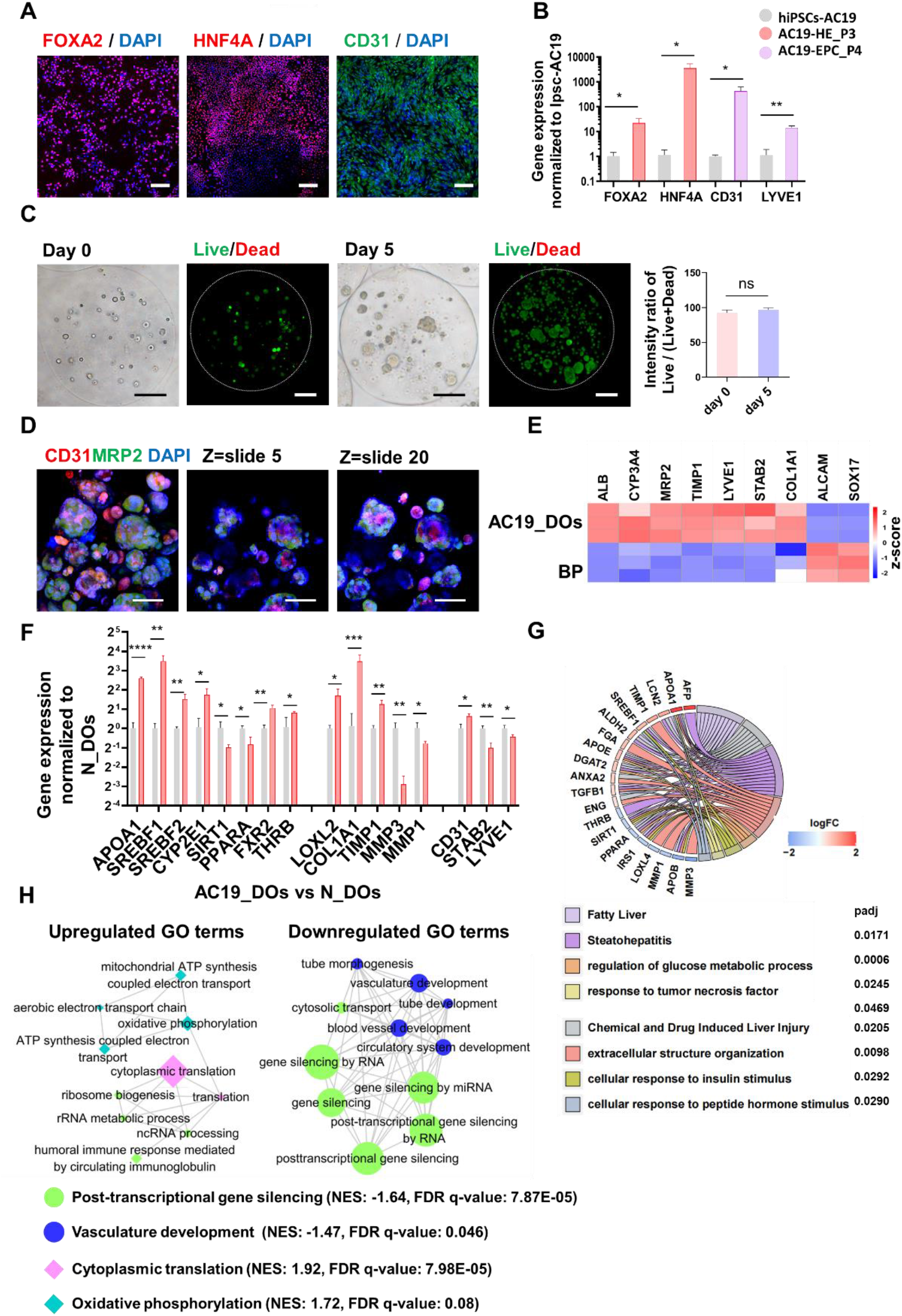
AC19-DOs have ALD transcriptional phenotypes. (A) Immunostaining of HE markers (FOXA2 and HNF4A) in hiPSCs-AC19_HE-P3 and endothelial marker (CD31) in hiPSCs-AC19_EPC-P4. Scale bars, 200 µm. (B) qPCR of the markers of HE and endothelial cells in hiPSCs-AC19, hiPSCs-AC19_HE-P3, and hiPSCs-AC19_EPC-P4, normalized to GAPDH and hiPSCs. (C) Optical images and maximal projection of live/dead fluorescent images of postprinted cell-laden microspheres at day 0 and day 5. Scale bars, 100 µm. (D) Maximal projection of CD31/MARP2/DAPI staining in AC19_DOs harvested from microspheres. The right two panels are slides 5 and 20. Scale bar, 50 µm. (E) Heatmaps of gene expression in AC19_DOs vs. BP; the bar stands for z score values calculated based on FPKM, n=3. (F) qPCR of ALD pathological hallmark genes in N_DOs and AC19_DOs normalized to GAPDH and N_DOs. In (B), (C), and (F), the data are presented as the mean ± s.d.; statistical significance was calculated by Student’s T test. \**p*<0. 05, \*\**p*<0. 01, \*\*\**p*<0. 001, \*\*\*\**p*<0. 0001, n=3. (G) Chord plot illustrating the relationships between ALD-related DEGs (left) and the terms enriched by DEGs of AC19_DOs vs. N_DOs (right). The bar represents the changed folds (log10 scale) of DEGs. (H) PPI network of representative GO terms enriched by GSEA. Circle nodes represent terms, and the size is inversely proportional to FDR q values (all <0. 25). The colors represent cluster identities.

Patient hiPSCs carry a personalized genetic and epigenetic background^24^. To investigate whether AC19_DOs represented ALD susceptibility, we performed qRT–PCR to compare the transcription of ALD biomarkers between AC19_DOs and N_DOs without exogenous stimulation (Fig. 5F). Compared to N_DOs, the transcription of biomarkers in AC19_DOs showed a pathologically relevant tendency in ALD^25-27^. The downregulation of LSEC positive (STAB2, LYVE1) and upregulation of negative (CD31) markers suggested the pathological gene-phenotype of LSECs^28^. RNA-seq analysis revealed that these differentiated biomarkers were enriched in metabolic liver disease-related terms (Fig. 5G). Noticeably, the enrichment of extracellular structure organization suggested that the 3D structure contributed to the replication of transcriptional characteristics of ALD (Fig. 5G).

To further reveal the representation of AC19_DOs on ALD inheritance, we conducted GSEA and PPI network analysis of GO terms between AC19_DOs and N_DOs. Cytoplasmic translation and mitochondrial oxidative phosphorylation were upregulated, consistent with the fact that mitochondrial stress is the hallmark of ALD pathology^25^. Posttranscriptional gene silencing and vasculature development were significantly downregulated and closely interacted (Fig. 5H). These results exemplify that alcohol led to the upregulation of NADPH oxidase-related genes in rat LSECs in vivo and human endothelial cells in vitro by decreasing miRNA^29^.

Similarly, nonalcoholic steatohepatitis (NASH) patients’ hiPSC-derived hepatocyte organoids exhibited higher lipid accumulation, a hallmark of NASH, than those from healthy controls in the absence of inducer^24^. We upgraded patient hiPSC-derived organoids from single to multicellular compositions and from naked to hydrogel scaffold-encapsulated, facilitating recapitulation of ALD pathophysiological transcriptional profiling without exogenous stimulation.

### 6. MP-TBBPA complexes exhibit hepatotoxicity to ALD patient-derived DOs

Five of six patients whose liver tissues accumulated MPs suffered ALD^3^. Given the accumulation in cirrhotic liver, the effect of MPs in liver disease remains elusive, i.e., whether they are a contributor or a consequence. In previous tests with N_DOs, MPTA upregulated the expression of IGFBP1, a diagnostic biomarker of ALD^30^. Here, to reveal the effect on the livers of ALD susceptible individuals, we investigated the cytotoxicity, transcriptomic and functional toxicities of MPTA on ALD patient-derived AC19_DOs at 600 ng/mL/100 nM with 3-day repeated exposure (Fig. 6A). Furthermore, we analyzed the interplay between MPTA and the transcriptional expression of ALD inheritance genes.

**Fig 6.**
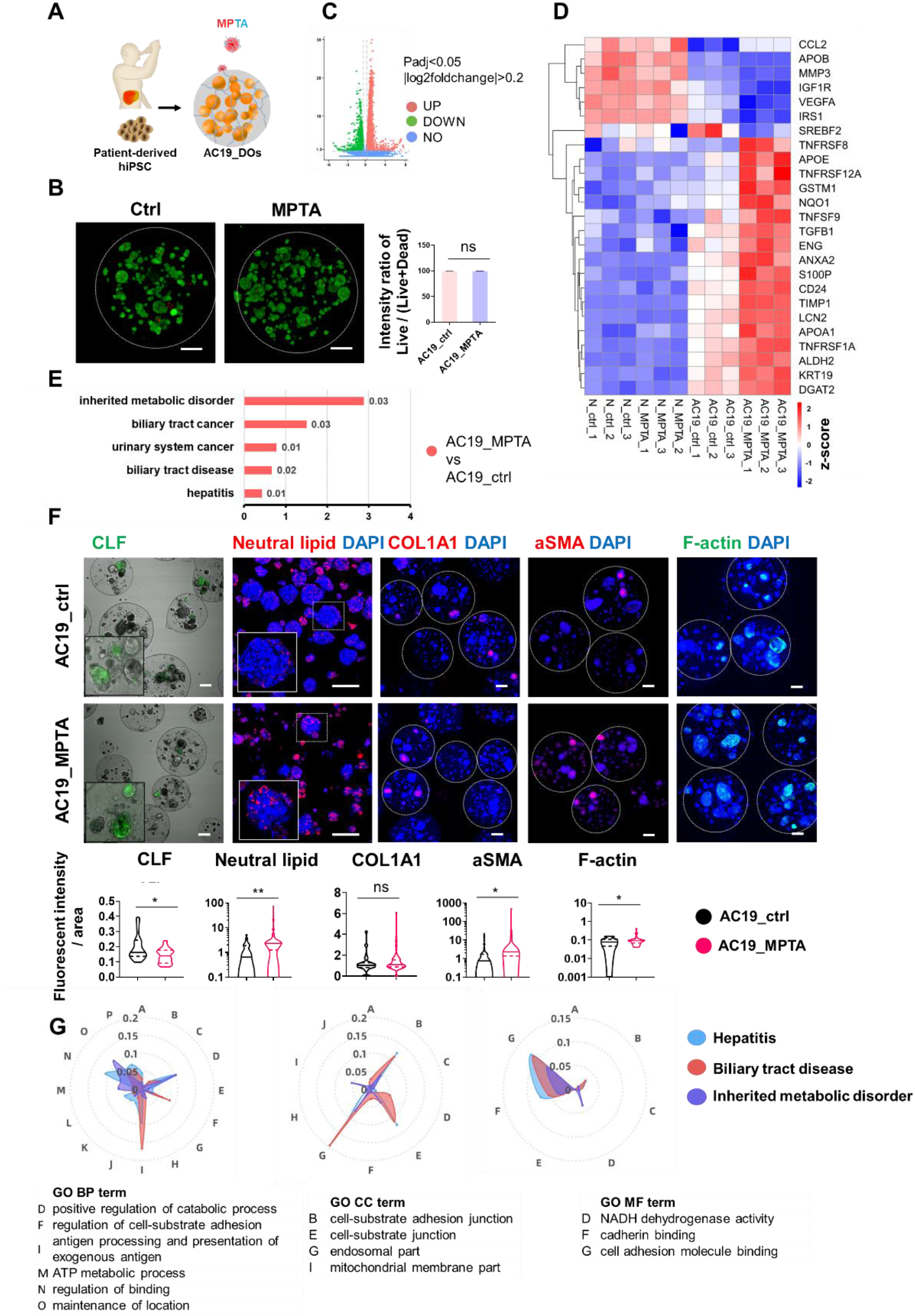
Hepatotoxicity of MPTA on AC19_DOs. (A) Schematic of the toxicity assessment of MPTA with AC19_DOs. (B) The volcano plot depicts the DEGs of AC19_MPTA vs. AC19_ctrl. (C) Heatmaps of ALD-related gene expression in N_DOs and AC19_DOs treated with vehicle (Ctrl) and MPTA; the bar represents z score values calculated based on FPKM, n=3. (D) Maximal projection of the in situ staining of Live/Dead in N_DOs within microspheres. Scale bars, 100 µm. The dashed circles mark the outline of the microspheres. (E) GSEA of DisGeNET terms based on the differential expression between AC19_DOs and N_DOs. The numbers are FDR q-values. (F) Neutral lipid staining in harvested AC19_DOs, in situ staining of CLF (slides), COL1A1 (the maximal projection), αSMA (the maximal projection) and F-actin (the maximal projection) in AC19_DOs within microspheres, with treatments of vehicle and MPTA. Scale bars, 100 µm. The lower panels are the quantification based on the staining images of CLF, COL1A1, aSMA, and F-actin. The statistical significance was determined by Student’s t test. \**p*<0.05, \*\**p*<0.01. (G) The radar chart plots the similarity scores between the GO terms and disease-related terms of hepatitis, biliary tract disease, and inherited metabolic disorder. The similarity scores are calculated based on shared gene numbers.

MPTA did not impair cell viability (Fig. 6B), consistent with the results obtained with N_DOs (Fig. 4B). RNA-seq analysis revealed the significant differentiation of transcriptome correlation between AC19_MPTA and AC19_ctrl (Fig. S4A, B). MPTA upregulated 2865 genes and downregulated 2122 genes (Fig. 6C, Fig. S4C), which were involved in the ribosome, mitochondrial functions, antioxidant activity, cell-substrate junction, and extracellular matrix components (Fig. S4D-E). MPTA disrupted ALD genetic transcription of AC19_DOs in a pathologically relevant tendency (AC19_MPTA vs. AC19_ctrl, Fig. 6D) despite not disrupting their expression in N_DOs (N_MPTA vs. N_ctrl, Fig. 3C). Hepatic lipid metabolism-related genes (SREBF1 and CYP2E1) were enhanced. Therefore, MPTA pathologically regulated the expression of ALD susceptibility genes.

To predict the disease that MPTA causes, we clustered the DEGs between AC19 MPTA and AC19 ctrl into the disease-specific database DisGeNET. Diseases at high risk include inherited metabolic disease, biliary tract cancer, urinary system cancer, biliary tract disease, and hepatitis (Fig. 6E). Functional analysis by immunostaining showed that MPTA impaired bile acid transport, aggravated neutral lipid accumulation and led to hepatic fibrosis associated with upregulation of αSMA and F-actin despite the lack of alteration of collagen I (Fig. 6F). This finding is consistent with the findings of elevated aSMA by MPs in rats^31^ and deteriorated lipid metabolism-related oxidative stress in the liver of zebrafish by MPs and TBBPA^6^.

Mechanistically, liver disease-related terms by MPTA were closely correlated with the cell-substrate adhesion junctions, cadherin binding and endosomal parts (Fig. 6G, S4D). This speculated mechanism was exemplified by the in vivo and in vitro findings that 1 µm microplastics affected adhesion protein expression in endothelial cells and heterogeneous cell adhesion^32^. Specifically, the inherited metabolic disorder that MPTA upregulated was correlated with mitochondrial membrane components (Fig. 6G, GO CC) and NADH dehydrogenase activity (Fig. 6G, GO MF), both of which are hallmarks of metabolic liver diseases^33^. Therefore, these results demonstrated that MPTA impaired bile acid transport, disrupted lipid metabolism, and led to hepatic fibrosis by disrupting cell‒cell/matrix interactions, endosome transport, and mitochondrial functions.

Transcriptome comparison of AC19_MPTA and AC19_ctrl represented the transcriptomic toxicity of MPTA (Group MPTA); AC19_DOs against N_DOs represented the ALD genetic suspensibility (Group models) (Fig. S4F). To reveal the interplay of ALD genetic suspensibility and MPTA, we analyzed the similarity of terms between Group MPTA and Group models. Approximately 80% of DEGs in the terms of Group models overlapped with Group MPTA (Fig. S4G and S4H), indicating that ALD genetics contributes to susceptibility to MPTA hepatotoxicity.

Taken together, MPTA caused hepatotoxicity in AC19_DOs, not N_DOs, suggesting that MPTA contributes to liver diseases specifically in ALD-susceptible individuals. These results exemplified that the interaction of hereditary factors and pollutants contributes to suspensibility to environmental toxicants^34,35^. MPs can transport organic pollutants into the body by adsorption. Our study explains co-exposure of MPs and organic pollutants as contributors to liver fibrosis, i.e., MPTA contributes to or worsens the pathogenesis of liver fibrosis in disease-susceptible populations by interfering with mitochondrial functions, cell‒cell/matrix interactions, and endocytic transport, which in turn increases the accumulation of MPs in the liver.

### 7. Conclusion

Tons of environmental contaminants, such as MPs detected in human liver tissues, need *in vitro* models to evaluate the toxicities^1,36^. Guided by the 3R principles (replacement, reduction, refinement), human organoids are a reliable alternative to animals in toxicology. High cell viability, biomimetic structure and functions, and reproducibly massive production are the key factors in the application of organoids in environmental toxicology.

We fabricated multicellular organoids (DOs) from monodispersed hiPSC-derived cells. Compared to prevailing 3D extrusion printing, microsphere printing technology is more suitable for the fabrication of hiPSC-derived multicellular organoids, with improved cell viability and more efficient cell assembly^8,12^. DOs resembled the close heterogeneous cell‒cell interaction in liver Disse space, Disse space-specific cell phenotypes, and multifaceted liver functions. They were more representative of the liver than PHHs in terms of endothelial functions, cell-ECM interactions, and hormone responses. DOs-laden microspheres can be regarded as independent mini liver units. The massive production, 10^7^ microspheres biofabricated from 1 mL bioink, facilitated applications for high-throughput screening of hepatotoxic contaminants.

For carrying inheritance information, hiPSCs have been proven effective for modeling monogenic mutant diseases in the context of differentiated monotypic cells^24,37^. This study found that ALD patient hiPSC-derived multicellular DOs can recapitulate ALD transcriptional profiling without inducers, suggesting the applicability of hiPSC-derived multicellular organoids in modeling diseases with complex regulatory mechanisms.

The hiPSC-derived DOs allowed us to evaluate the hepatotoxicity of environmental toxicants in the context of individual inheritance. MPs were detected in human cirrhosis liver tissues^3^. In the DOs, MPs not only accumulated but were also internalized by cells and appeared in the nuclear region. TBBPA, which is metabolized in the liver, promoted MP diffusion, highlighting the synergistic toxic effects. Environmental dosages of MPs and TBBPA indeed impacted the majority of gene transcription and liver functions, intriguingly, in ALD patient-derived DOs, not healthy individual-derived DOs. These results were associated with the detection of MPs in cirrhotic livers but not in healthy livers. These results support the views that pollutants and inherited disease genes contribute to susceptibility to toxicant toxicity and liver diseases^38^. MPs and TBBPA are severely toxic to aquatic organisms, especially in hepatotoxicity^4,39^. We presumed that disease-susceptible liver and aquatic organisms share some genes that make them susceptible to the toxicity of contaminants such as MPs and TBBPA.

Our study calls for attention to the individual differences in the toxic effects of some contaminants, and we should pay more attention to the toxicity of the pollutant on disease-susceptible people. This study provides a proof of principle for applying biofabricated hiPSC-derived multicellular liver organoids in personalized environmental toxicology.

## Materials and Methods

### 1. Cell culture

hiPSCs-N were reprogrammed from the peripheral blood cells of a healthy male (Nuwacell, RC01001-A). hiPSCs-AC19 were reprogrammed from the peripheral blood cells of a 64-year-old male patient with ALD. The patient provided written consent, and an ethical statement was obtained from the Institution Review Board of Tsinghua University. The reprogramming of hiPSCs-AC19 was performed by the Nuwacell company. Both hiPSCs expressed pluripotent stem cell markers, maintained normal karyotypes, and formed teratomas in vivo. hiPSCs were cultured in Matrigel (BD, 354277)-coated tissue culture plates in ncTarget medium (Nuwacell, RP01020). The medium was refreshed daily. The hiPSCs were split at a ratio of 1:10 when they reached 80% confluence. LX-2 cells (Procell, CL-0560) were cultured in DMEM (Gibco, 11965175) supplemented with 5% FBS and 1% antibodies. The culture medium was refreshed every two days.

### 2. Differentiation of hiPSCs into HE and EPC

#### 2.1 HE specification and expansion

hiPSCs were dissociated into single cells by Nuwacell® Solase (RP01021), and 1×10^5^/mL cells were seeded onto Matrigel (BD, 354230)-coated 12-well plates with ncTarget medium plus 10 µM Y27632 (Selleck, S6390). The media recipes are shown as follows. On days 0-1, RPMI 1640 (Thermo Fisher, 11875119) was supplemented with 1×B27 insulin minus (Gibco, A1895601) and 3 µM CHIR99021 (Selleck, S1263). On days 2-3, the medium was supplemented with 100 ng/mL activin A (R&D, 338-AC). On days 4-7, the optimized HE induction medium included RPMI 1640, 1×B27 (Gibco, A1486701), 20 ng/mL BMP4 (PeproTech, 96-120), and 10 ng/mL FGF2 (PeproTech, 96-100-18B). HE cells were obtained on day 8.

On day 8, the cells were counted as HE_P0. Then, the HE_P0 cells were dissociated and split at a ratio of 1:3 in a defined medium composed of DMEM-F12 (Gibco, 11320033), 1× ITS (Merck, I3146), 500 µM monothioglycerol (Sigma–Aldrich, M6145), 1× GlutaMax (Gibco, 35050061), 50 µg/mL ascorbic acid (Sigma–Aldrich, A8960), 0.1% BSA (Sigma–Aldrich, V900933), 5 ng/mL FGF2, 10 ng/mL VEGF (R&D Systems, 293-VE), 20 ng/mL EGF (PeproTech, AF-100), 3 µM CHIR99021, and 5 µM A83-01 (Selleck, S7692). The medium was refreshed every day.

#### 2.2 EPC specification and expansion

EPCs were derived from hiPSCs according to the protocol^40^. Briefly, hiPSCs at 4×10^5^/mL were seeded onto Matrigel-coated 12-well plates and cultured overnight in ncTarget medium supplemented with 1 µM CHIR99021 and 10 µM Y27632. The media recipes are shown as follows. On days 0-1, RPMI 1640 was supplemented with 1× B27 insulin minus, 50 ng/mL Activin A and 1/60 Matrigel (BD, Growth Factor Reduced, 354230). On days 1-2, RPMI 1640 was supplemented with 1× B27 insulin minus, 40 ng/mL BMP4, and 1 µM CHIR99021. On days 2-5, the optimized basal medium included Vivo15 (Lonza X-VIVO15, 04-418Q), 1× ITS, 500 µM monothioglycerol, 1× GlutaMax, 50 µg/mL ascorbic acid, and 0.1% BSA; the optimized basal medium was supplemented with 300 ng/mL VEGF, 5 ng/mL FGF2 and 10 ng/mL BMP4. On day 5, the cells were obtained and continuously cultured in optimized basal medium supplemented with 20 ng/mL FGF2, 20 ng/mL VEGF, and 1 µM CHIR99021. When confluent, the cells were counted as EPC_P0 and split at a ratio of 1:3. The medium was refreshed daily.

### 3. Fabrication of DO-laden microspheres by electroassisted inkjet printing

The printing procedure refers to Yao’s paper^41^. The bioink was composed of 3.5% alginate (Sigma, 919373), 2 µg/mL ncLaminin 511 (Nuwacell, RP01025), and a cell mixture of hPSC-HE, hiPSC-EPCs, and LX-2 at 10:7:2 and a total density of 3×10^6^ cells/mL.

The electroassisted inkjet printing device consists of a static electricity power supply, a syringe pump (Longer Pump Ltd.), and a grounded collecting device. A disposable sterile syringe loaded with bioink was fixed to the syringe pump. Cell-laden microspheres were printed at a 10 mL/h propulsion speed, 2 cm electrode distance, and 12 kV voltage. Microspheres were cross-linked with 300 mM CaCl_2_ solution for one minute and washed with DMEM-F12 basal medium three times.

The postprinted cell-laden microspheres were cultured in organoid culture medium composed of DMEM-F12/M199/William (1:1:1), 500 µM monothioglycerol, 1×B27 with insulin, 1% ITS, 50 µg/mL ascorbic acid, 0.1% BSA, 2% FBS, 1% v/v sodium pyruvate (Sigma, P2256), 1% v/v NEAA (Sigma, M7145), 1% v/v GlutaMAX, 100 µM ascorbic acid, 20 ng/mL VEGF, 20 ng/mL HGF (R&D, 294-HG-025), and 5 ng/mL FGF2. The medium was refreshed daily. DOs-laden microspheres were formed after 5 days of culture.

### 4. Pollution exposure

The spherical polystyrene MPs (Wuxi Ruige Biotechnology Co., LTD) are monodispersed particles with a diameter of 1 µm in water at a concentration of 25 mg/mL. The stock solutions were pasteurized before utilization. TBBPA (purity > 98.0%, TCI, Shanghai Development Co., Ltd., China) was dissolved in dimethyl sulfoxide (DMSO, Amresco, USA, 0231). Polystyrene MPs were diluted in sterile water; TBBPA was diluted in DMSO. The MPTA complex was obtained by incubating 6 µg/mL PS-MPs and 10 µM TBBPA in medium at 37 °C overnight.

The exposure dosage of PS-MPs was 600 ng/mL, TBBPA was 100 nM, and MPTA was 600 ng/mL/100 nM. The vehicle control was 0.01% DMSO. DOs-laden microspheres were incubated with MPs, TBBPA, MPTA, and vehicle control for 3 days. The media were changed every day.

### 5. Tracing MPs in DOs

The cell membranes of hiPSC-derived HE, EPC, and LX-2 cells were labeled with a PKH67 Green Fluorescent Cell Linker Kit (Merck, PKH67GL) according to the manufacturer’s protocols. We printed three types of microspheres with cell mixtures: PKH67-labeled HE, unlabeled EPC, and LX-2; PKH67-labeled EPC, unlabeled HE, and LX-2; PKH67-labeled LX-2, unlabeled HE, and EPC. After 5 days of coculture, the three types of DO-laden microspheres were obtained.

To observe the process of microplastic entry into cells more clearly in a short period, we used higher dosages of MPs than environmental concentrations. The DO-laden microspheres were exposed to 100 µg/mL red fluorescence-labeled MPs (for better visualization) (Wuxi Ruige Biotechnology Co., LTD) and MPTA at 100 µg/mL/100 nM. The DO-laden microspheres were placed in a sterile chamber at 5% CO_2_ and 37 °C.

The diffusion of MPs in the DOs was visualized and recorded by a confocal microscope (LSCM, Nikon, Z2). The fluorescence signal distribution of MPs and PKH67 at the specified locations was analyzed and graphed by Nikon NIS-Elements AR. The fluorescence intensities and areas of the more than 10 microspheres were analyzed by Nikon NIS-Elements AR. The fluorescence intensity of the MPs (n>10) was the average of the normalized fluorescence intensity of MPs, which was divided by the area of the microsphere.

### 6. Cell live/dead staining

The cell viability in the postprinted microspheres, DOs within microspheres, and DOs with pollutant treatments was analyzed by in situ staining of microspheres for live/dead cells. The live/dead cells were stained with the Calcein-AM/PI Kit (Merck, 04511-1KT-F) according to the manufacturer’s instructions. The staining was incubated for 10 min in the dark and observed using a confocal microscope (LSCM, Nikon, Z2). Cell viabilities were the averages of the ratio of green fluorescence intensity to total cell fluorescence intensity, n>3.

### 7. Immunofluorescence staining

#### 7.1 Staining of cultured planner cells

hiPSC-derived HE and EPC were fixed with 4% paraformaldehyde for 15 min and blocked with DPBS (Solarbio, D1040) containing 5% goat serum (Solarbio, China, SL038) and 0.1% Triton X-100 (Amresco, USA, 0694-1 L) for 1 hour at room temperature. Then, the cells were incubated with primary antibodies for 1 hour and secondary antibodies for 30 min at room temperature.

#### 7.2 Staining of DOs

The DO-laden microspheres were incubated in a decrosslinking solution (150 mM sodium chloride containing 55 mM sodium citrate and 20 mM EDTA), and DOs were harvested by gravitational settlement.

The harvested DOs or DO-laden microspheres were fixed with 4% paraformaldehyde for 45 min at 4 °C, washed three times with 0.1% (vol/vol) Hanks (Solarbio, H1025)-Tween-20, and blocked with Hanks containing 2% BSA and 0.1% Triton X-100 for 15 min at 4 °C. Then, the samples were incubated with primary antibodies overnight at 4 °C. After being washed three times, the DOs were continuously incubated with secondary antibodies overnight at 4 °C. Fluorescent images were taken by confocal microscopy (LSCM, Nikon, Z2).

The fluorescent intensities of COL1A1, αSMA and F-actin were analyzed by Nikon NIS-Elements AR. The fluorescence intensity in each slide was normalized to the DAPI area. The fluorescence intensity of each target protein was calculated by the averages of all the slides in three independent experiments.

The following antibodies and dilution ratios were employed: HNF4α (1:200, Abcam, ab92378), FOXA2 (1:200, Abcam, ab256493), ALB (1:100, Abcam, ab83465), CD31 (1:100, Cell Signaling Technology, 89C2), MRP2 (1:100, Abcam, ab3373), αSMA (1:100), COL1A1 (1:100, Cell Signaling Technology, 72026), F-actin (Invitrogen, A12379), Alexa Fluor® 594-conjugated secondary antibody (1:1000, Abcam, ab150080) and Alexa Fluor® 488-conjugated secondary antibody (1:1000, Abcam, ab150113). The nuclei were stained with DAPI (Sigma, D9542).

### 8. Liver function detection

#### 8.1 Chyl-lysyl-fluorescein (CLF) staining

Bile salt transport was checked by chyl-lysyl-fluorescein (CLF) staining. DOs-laden microspheres were incubated with Williams’ E medium (BasalMedia, L660KJ) plus 5 µM cholyl-lysyl-fluorescein (CLF, BD-451041) for 15 minutes at 37 °C and 5% CO_2_. After washing, CLF internalization was visualized with confocal microscopy. The fluorescence intensity of CLF was analyzed by Nikon NIS-Elements AR. More than 20 DO-laden microspheres from three independent experiments were included in the calculation. The CLF intensities were calculated as the average fluorescence intensity of the CLF divided by the microsphere area.

#### 8.2 Neutral lipid staining

Lipid accumulation was detected by neutral lipid staining. DOs were harvested from microspheres, fixed in 4% formaldehyde for 30 min, washed and stained with red neutral lipid dye (Invitrogen, H34476) according to the manufacturer’s instructions. The nucleus was stained with DAPI. The DOs were visualized with confocal microscopy. The fluorescence intensity of neutral lipids was analyzed by Nikon NIS-Elements AR. The fluorescence intensity of neutral lipid dye in each slide was normalized to the DAPI area. Then, the average of all the slides in three independent experiments was counted as the fluorescence intensity of neutral lipid staining.

#### 8.3 Indocyanine green (ICG) uptake assay

The indocyanine green (ICG) uptake assay was performed by adding 1 mg/mL ICG to the medium for 30 min. ICG was wholly released from the cells after 7 hours. ICG uptake and release images were captured from more than 20 microspheres using an inverted optical microscope. Image-Pro software was used to calculate the uptake ratio as the IOD of the microsphere divided by the area of the microsphere.

### 9. Total RNA isolation and qRT–PCR analysis

The DOs were harvested from microspheres with a decrosslinking solution. The DOs were resolved in TRIzol (Life Technologies, USA, 15596018). Total RNA samples were isolated according to the TRIzol manufacturers’ specifications. RNA (2 µg) was reverse transcribed into cDNA with the RT Reagent Kit with gDNA Eraser (Takara, Japan, RR047A). qPCR was performed using a SYBR Premix Ex Taq kit (Takara, Japan, RR420A) on a QuantStudio 6 Flex System (Thermo, USA). The PCR primers are listed in Table S1.

### 10. RNA-seq

The samples for RNA-seq were collected in TRIzol. RNA isolation, cDNA library construction, sequencing and primary analysis were performed at Novogene (Tianjin, China). Briefly, 1 µg of RNA per sample was used as input material for the RNA sample preparations. Following the manufacturer’s recommendations, sequencing libraries were generated using the NEBNext® UltraTM RNA Library Prep Kit for Illumina® (NEB, USA), and index codes were added to attribute sequences to each sample. RNA quality was analyzed with an Agilent 2100 Bioanalyzer and reached the standard.

Sequencing was performed with an Illumina NovaSeq platform, and 150 bp paired-end reads were generated. Reads were filtered and mapped to the reference genome sequence with HISAT2 v2.0.5. Feature Counts v1.5.0-p3 was used to count the read numbers mapped to each gene. The fragments per kilobase of exon per million fragments mapped (FPKM) method was used to calculate gene expression levels. Differentially expressed genes (DEGs) with fold change ≥ 1.25 and FDR q-value <0.05 were calculated using the DESeq2 R package (1.16.1).

GO, KEGG, Reactome and DisGeNET enrichment analyses of DEGs were implemented by the clusterProfiler R package. Significantly enriched GO and KEGG terms were determined based on a corrected p value <0.05. Based on KEGG, Reactome, and DisGeNET datasets, GSEA was conducted by the local version of the GSEA tool http://www.broadinstitute.org/gsea/index.jsp. A PPI network of representative GSEA GO terms from the whole clusters was constructed with Cytoscape EnrichmentMap. The Pearson correlation heatmap, volcanic map and gene heatmap were calculated and graphed by R (Version 3.0.3) ggplot2 and pheatmap packages. The chord plot was graphed by R GOplot v1.0.2. R (Version 3.0.3). The parallel set was graphed by Origin 2021 software. All original sequence datasets were submitted to the GEO database under the accession **number GSE212099**.

### 11. Statistical analysis

Data are expressed as the means ± SD and were analyzed using Prism software (GraphPad Inc.). Statistical analysis was performed using Student’s t test or one-way ANOVA. A statistically significant difference and replicates are shown in the legends.

## Acknowledgments

The authors are sincerely grateful for funding from the National Key Research and Development Program of China (2018YFA0109000) and the National Natural Science Foundation of China (31871015, 31927801, and 61675059).

## Author contributions

Rui Yao and Shaojun Liang conceived the project, contributed to RNA-seq and data analysis, and prepared the figures. Shaojun Liang and Yixue Luo fabricated the DO-laden hydrogel microspheres and performed the experiments. Shaojun Liang, Lu Feng, and Yijun Su contributed to the immunostaining of organoids, measurement, and quantitative analysis. Shaojun Liang and Yixue Luo wrote the Methods and Supplementary section. Shaojun Liang, Rui Yao and Mingen Xu wrote and revised the manuscript.

## Data availability

The bulk RNA-seq data were submitted to the GEO database (GEO GSE212099).

---

**Fig. S1.**
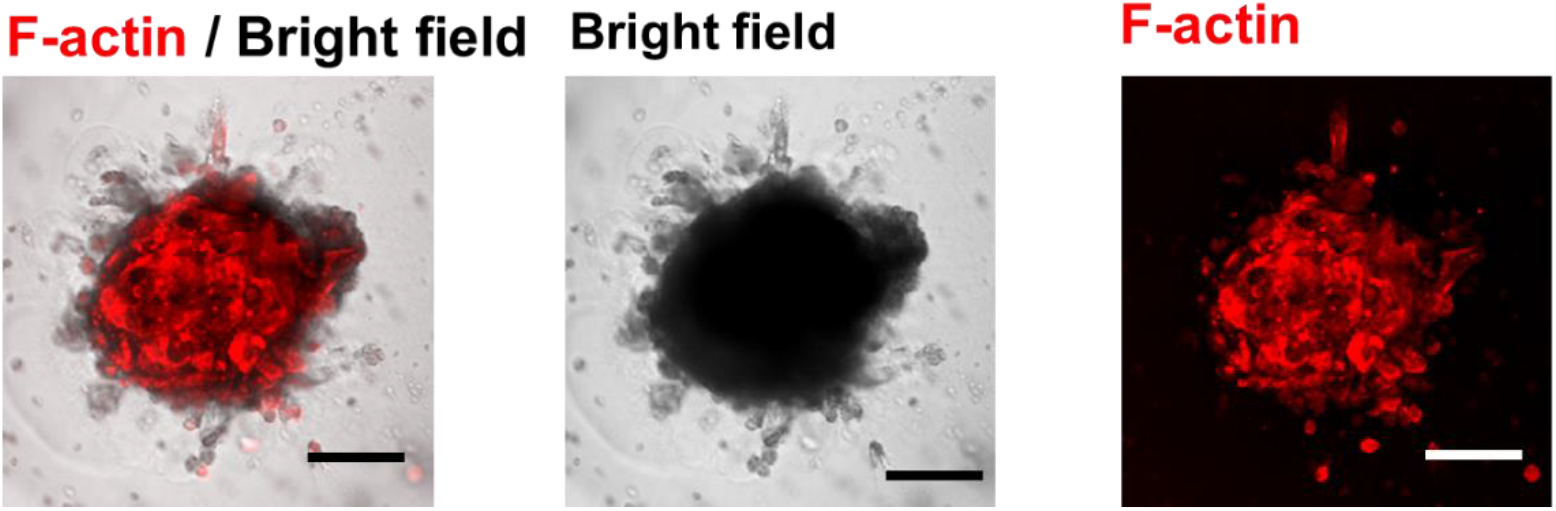

**Fig. S2.**
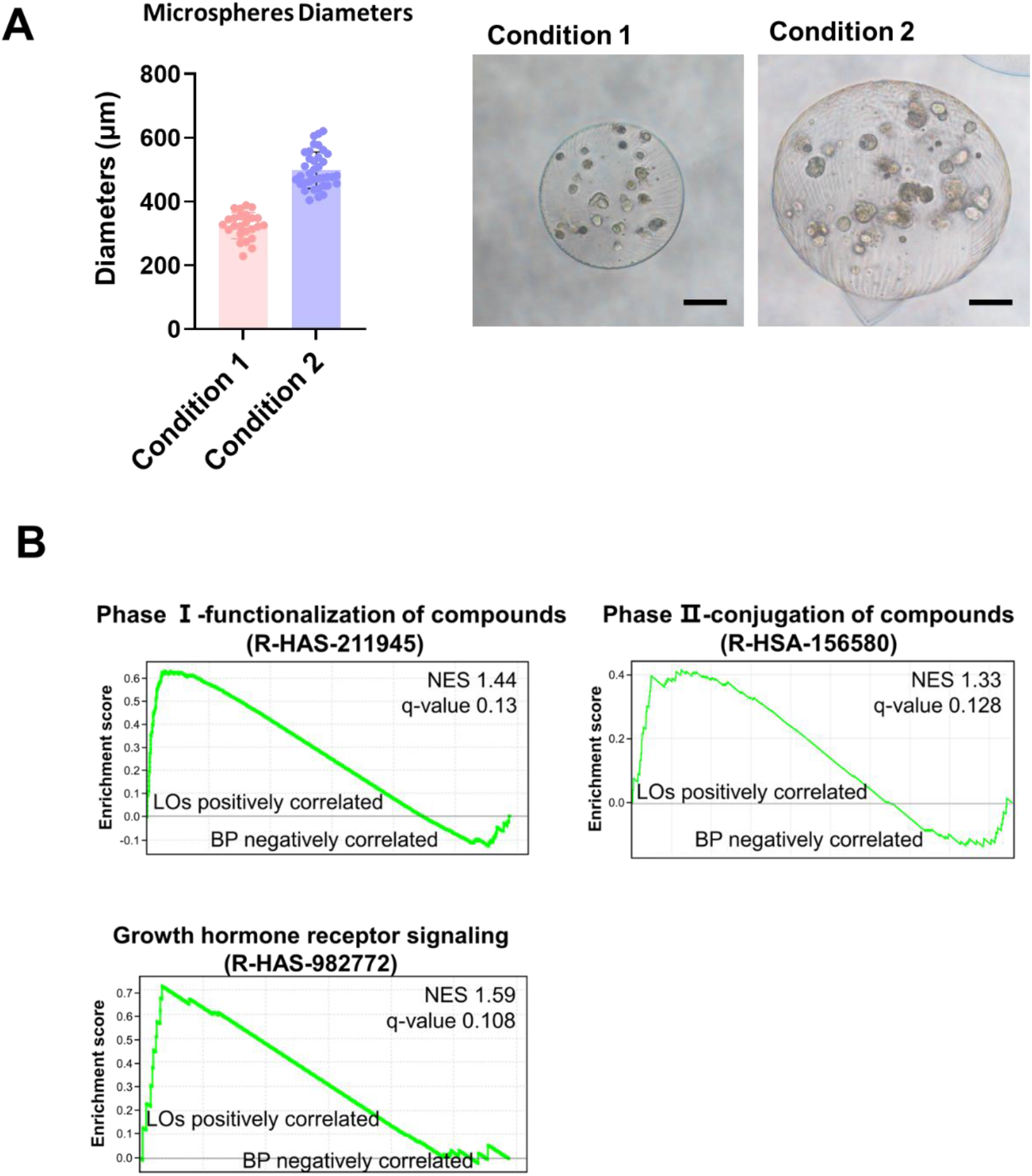

**Fig. S3.**
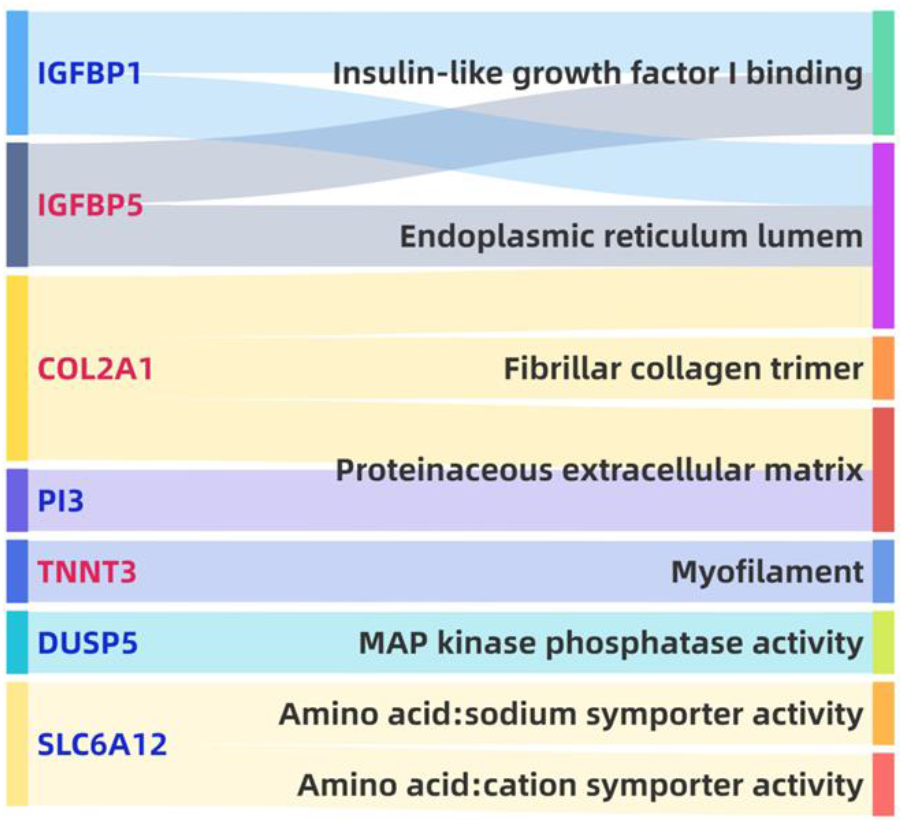

**Fig S6.**
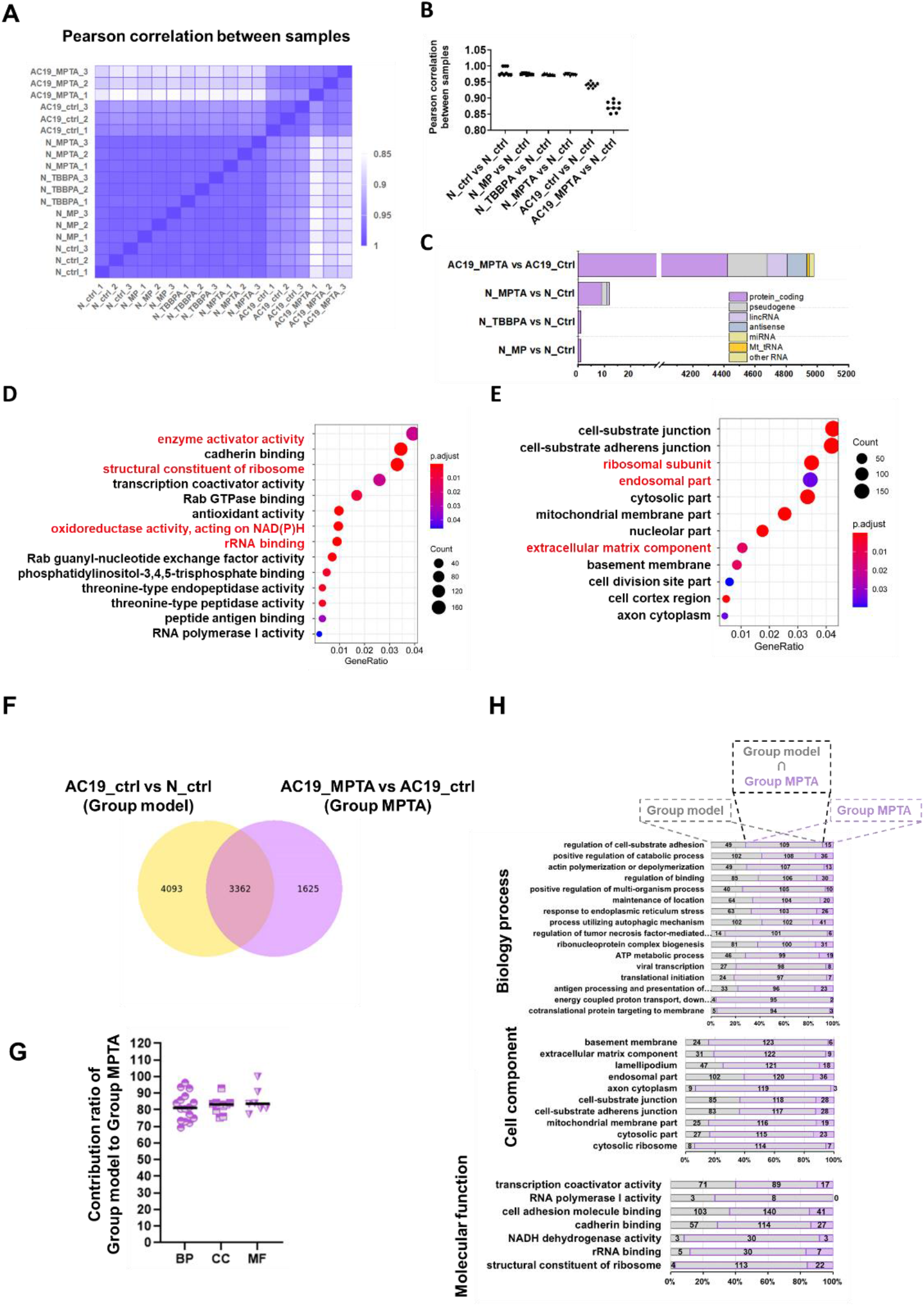
Interplay of MPTA and ALD susceptible genes of AC19_DOs. Heatmap (A) and plot (B) of correlation analysis of the gene counts of the whole transcriptome of AC19_MPTA, AC19_ctrl, N_MPTA, N_TBBPA, and N_MPs, N_ctrl. n=3. (C) The DEGs numbers of AC19_MPTA compared to AC19_ctrl and N_MPTA, N_TBBPA, and N_MPs compared to N_ctrl, respectively. GO MF (D) and GO CC (E) enriched by the DEGs of AC19_MPTA vs. AC19_ctrl. (F) The venn of DEGs of group model and group MPTA. (H) The shared percentages of genes in key terms clusterd in Group model and Group MPTA. (G) The plot of contribution ratio of Group model to Group MPTA.

